# Dynamic DNA methylation turnover in gene bodies is associated with enhanced gene expression plasticity in plants

**DOI:** 10.1101/2022.12.02.518885

**Authors:** Clara J. Williams, Dawei Dai, Kevin A. Tran, J. Grey Monroe, Ben P Williams

## Abstract

**Background:** In several eukaryotes, DNA methylation occurs within the coding regions of many genes, termed gene body methylation (GbM). Whereas the role of DNA methylation on the silencing of transposons and repetitive DNA is well understood, gene body methylation is not associated with transcriptional repression, and its biological importance has remained unclear.

**Results:** We report a newly discovered type of GbM in plants, which is under constitutive addition and removal by dynamic methylation modifiers in all cells, including the germline. Methylation at Dynamic GbM genes is removed by the DRDD demethylation pathway and added by an unknown source of de novo methylation, most likely the maintenance methyltransferase MET1. We show that the Dynamic GbM state is present at homologous genes across divergent lineages spanning over 100 million years, indicating evolutionary conservation. We demonstrate that Dynamic GbM is tightly associated with the presence of a promoter or regulatory chromatin state within the gene body, in contrast to other gene body methylated genes. We find Dynamic GbM is associated with enhanced gene expression plasticity across development and diverse physiological conditions, whereas stably methylated GbM genes exhibit reduced plasticity. Dynamic GbM genes exhibit reduced dynamic range in *drdd* mutants, indicating a causal link between DNA demethylation and enhanced gene expression plasticity.

**Conclusions:** This study proposes a new model for GbM in regulating gene expression plasticity, including a newly discovered type of GbM in which increased gene expression plasticity is associated with the activity of DNA methylation writers and erasers and the enrichment of a regulatory chromatin state.

## BACKGROUND

In diverse eukaryote species, DNA methylation is important for transcriptional silencing of transposable elements [1,2], repressing recombination at repetitive regions [3] and establishing genomic imprinting [4,5]. DNA methylation is a reversible and dynamic mark, which can be added or removed by a number of writer or eraser enzymes [6]. Many of these writer pathways have evolved to promote highly stable inheritance of DNA methylation over cell divisions and reproductive generations [7,8]. For example, the enzyme MET1 (also known as DNMT1 in animals) acts on hemimethylated DNA at the replication fork, faithfully copying existing methylation patterns to the daughter strand at symmetrical CG sites [6]. Consequently, the vast majority of methylated CG sites are highly stable and consistently methylated across all cells [9]. In plants, many additional writers also maintain and methylate DNA de novo at CG, CHG and CHH sites, including the CHROMOMETHYLASE (CMT) family and the DOMAINS-REARRANGED METHYLTRANSFERASE (DRM) family [6]. Additionally, a family of plant-specific DNA glycosylase enzymes (collectively named DRDD enzymes) act as methylation erasers, removing DNA methylation from thousands of loci within the genome [10]. This family is comprised of the enzymes DEMETER (DME) [11], REPRESSOR OF SILENCING 1 (ROS1) [12], and DEMETER-LIKE2&3 (DML2/DML3) [13]. The DRDD pathway is critical for reproductive development [11,14,15], and also maintains epigenetic homeostasis by protecting transcribed genes from silencing by pathways targeting proximal repetitive DNA [12,16–20]. Additionally, the DRDD pathway can also act during somatic development to generate differences in the levels of DNA methylation between tissues [10].

These methylation writers and erasers together shape DNA methylation patterns across the genome, into two broad categories: 1) “heterochromatic” DNA methylation, in which cytosines are methylated in CG, CHG and CHH sequence contexts, abundant at repetitive DNA, heterochromatin and some intergenic sequences within euchromatin [6], and 2) “gene body” DNA methylation (GbM), in which the coding region of genes are methylated solely in the CG sequence context [9]. Whereas the function of heterochromatic DNA methylation is broadly understood, primarily functioning in the transcriptional silencing of repetitive DNA [1,21,22], or the potent regulation of certain specialized target genes [17,18,23,24], the functional role of GbM has remained more enigmatic [9,25,26]. GbM is associated with moderate-to-highly expressed genes [26,27], is inherited with high accuracy [28] and is conserved at homologous target genes across lineages [25]. However, a mechanistic link between GbM and gene regulation has been difficult to establish. Some lineages have dispensed with GbM altogether [9], raising questions about its necessity for gene regulation. Indeed, DNA methylation writer mutants or naturally-occurring genomes with reduced GbM do not display widespread changes to the transcriptional output of GbM genes, or strongly perturbed phenotypes [27,28]. Why GbM occurs in so many eukaryotic genomes, and at conserved genes in distinct lineages is therefore yet to be fully resolved.

The majority of GbM genes in the genome exhibit consistent methylation of CGs within all cells. In this study, we show that a subset of GbM genes exhibit heterogeneous methylation patterns between cells, indicating dynamic modification to methylation during development. This dynamic methylation heterogeneity is due to the activity of both writer and eraser pathways in all cell and tissue types, is evolutionarily conserved, and represents a newly discovered epigenetic state associated with regulatory histone modifications not typically found within gene bodies. Lastly, we show that Dynamic GbM genes exhibit dramatically higher gene expression plasticity than their stably methylated counterparts, suggesting that active modification to GbM may permit a greater exploration of the gene expression landscape.

## RESULTS

### DNA demethylation targets a subset of gene bodies

We recently generated a quadruple somatic mutant (*drdd*) of the four active DNA demethylases (DRDD enzymes) in the model plant *Arabidopsis thaliana* [10]. The global difference in CG methylation levels between WT and *drdd* mutants is small (23.9% vs 25.5%), but our previous study identified large effect methylation differences at thousands of individual loci throughout the genome. Unlike previously studied DNA demethylation mutants, *drdd* mutants exhibited increased CG methylation at a subset of gene bodies. To understand this genic demethylation further, we compared the 581 genes most evidently targeted by DRDD to previously described gene-body methylated (GbM) genes [9] (Fig. 1A). These genes were identified based on two cutoffs: the presence of a WT vs *drdd* differentially methylated region (DMR) within the gene body (see methods), and an additional filter requiring at least 5 CG dinucleotides (10 CG cytosines total) that gained >20% methylation in *drdd* mutants. We found that the genes targeted by DRDD were a mostly distinct group compared to previously identified GbM genes (Fig. 1B), exhibiting a number of different properties. Previously identified GbM genes exhibit high levels of CG methylation throughout the gene body (Fig. 1A, C), and each individual CG is consistently highly methylated across the majority of cells (Fig. 1A, E), much like the vast majority of stable across cells, we hereafter refer to these genes as Stable GbM genes. In contrast, genes targeted by DRDD display reduced methylation overall, in large part due to the fact that the majority of methylated CGs are methylated in only a *subset* of WT cells (in *drdd* mutants, these CGs are methylated in all cells, much like Stable GbM genes (Fig. 1A,E)). We refer to these genes as Dynamic GbM genes, due to the active removal of methylation that is occurring during plant development. The intermediate methylation state of CGs within Dynamic GbM genes (methylated in 10-85% of cells) is unusual and distinct from the vast majority of the genome (Fig. 1D-E), as the methylation maintenance by MET1 typically copies existing methylation states during DNA replication to ensure consistency across cell populations. A list of Dynamic and Stable GbM genes is available in Additional File 1.

**Figure 1:**
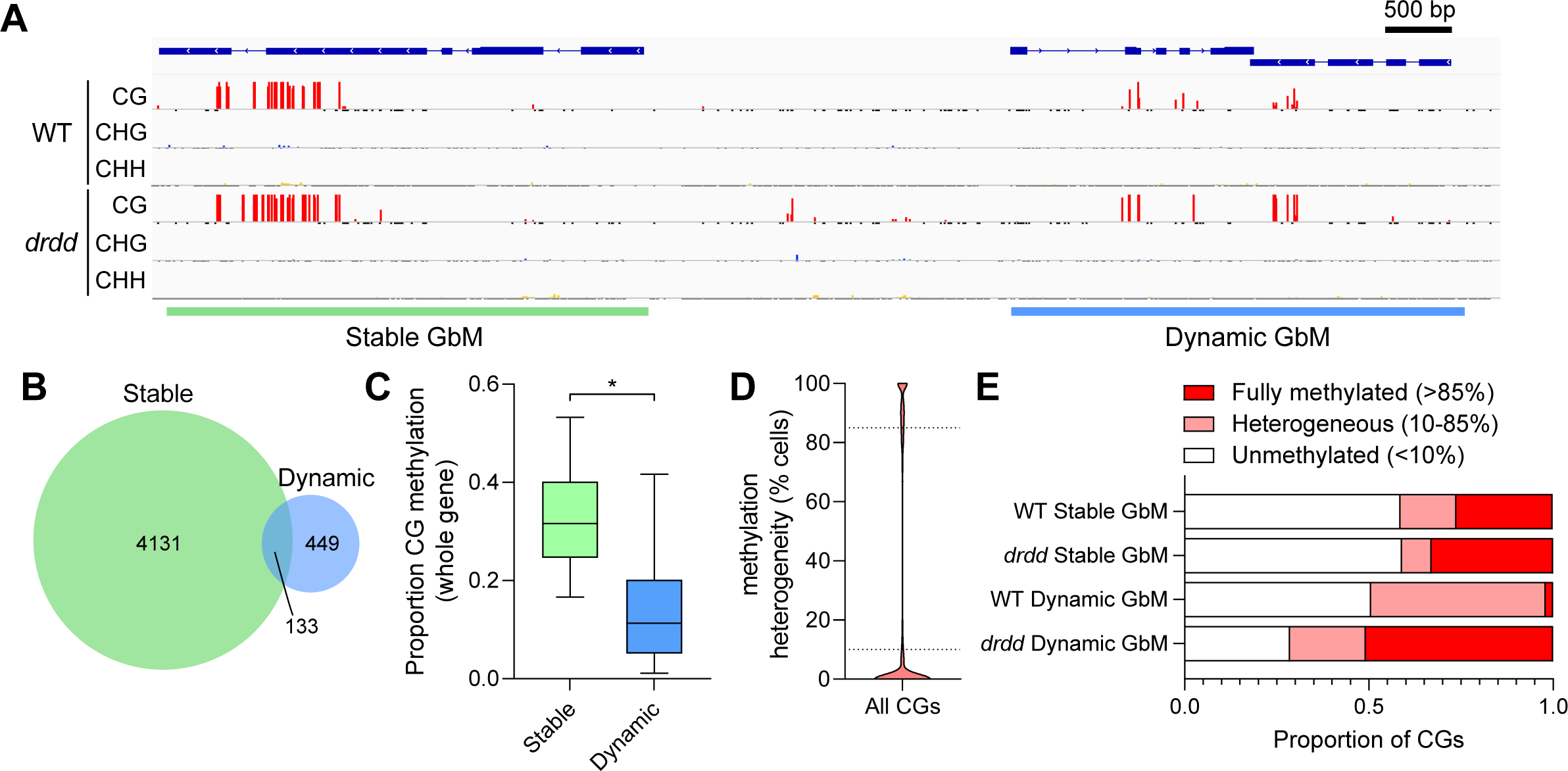
Two distinct types of Gene body methylation (GbM) in the Arabidopsis genome. A) Genome browser snapshot showing representative example of a gene with Stable GbM and a gene with Dynamic GbM in both WT and *drdd* mutants. Red bars represent methylated cytosines, and the height of the bar represents the percentage of cells in which a cytosine is methylated (scale = 0-100%). B) A Venn diagram showing the number of Stable and Dynamic GbM genes and the overlap between the two groups. C) Box plot showing the overall percentage CGs methylation across entire genes (all CGs across all cells) with Stable or Dynamic GbM. Line = median, box = interquartile range, whiskers = 5^th^ and 95^th^ percentile. *p = <0.0001 (two-tailed t-test). D) Violin plot showing the distribution of methylation heterogeneity levels of all CGs in the Arabidopsis genome, shown as the percentage of cells in which a CG is methylated. E) The proportion of fully methylated, heterogeneously methylated and unmethylated CGs within both Stable and Dynamic GbM gene bodies, in both WT and *drdd* mutants.

### DNA demethylation generates cellular heterogeneity in all tissues and cell types

As Dynamic GbM genes exhibit DNA demethylation in a subset of cells, we sought to determine the underlying developmental context of these methylation dynamics. It is possible that DRDD removes methylation in a specific subset of differentiated cell types within somatic tissues, or that the reduced methylation is developmentally specific to the tissue type (rosette leaves) of our sample. To test this, we assembled whole-genome methylation sequencing data from a number of studies (Additional file 2: Table S1) that have defined the methylation states of a number of cell and tissue types within Arabidopsis [10,29–32], and examined the cellular heterogeneity of DNA methylation patterns of both Stable GbM and Dynamic GbM genes. To our surprise, we discovered that the intermediate methylation of Dynamic GbM genes was not specific to active demethylation of a specific tissue or cell type (Fig. 2A-B) within somatic tissues, but rather a feature of all the tissue and cell types we examined. This suggested that the removal of methylation by DRDD is fairly ubiquitous across somatic development. Consistent with this, we did not find a relationship between the expression of DRDD genes and methylation heterogeneity of tissue types (Additional file 1: Fig. S1). By contrast, and as expected, Stable GbM genes were consistently methylated in the majority of somatic cells (Fig. 2B). As Dynamic GbM genes show cellular heterogeneity in DNA methylation patterns (a signature of active demethylation) in all tissues, we next wished to examine if the stem cell population in shoot meristems and germline cells also display cellular heterogeneity, or if the Dynamic GbM state is restricted to somatic development. By analyzing the cellular heterogeneity of CG methylation in shoot apical meristem stem cells (labelled with the *CLV3* promoter [32]), meiocyte cells from the male gametophyte [31] and sperm cells [30], we clearly observe intermediate methylation of CG sites across cells in all cases (Fig. 2C-D). To confirm that the cellular heterogeneity of CG methylation was due to the active removal by the DRDD pathway, we next examined DNA methylation in sperm nuclei mutant for one or both of the major DRDD genes *DME* and *ROS1* [30]. In demethylase mutant sperm, Dynamic GbM CG methylation patterns were homogenous across 100% of cells, conclusively demonstrating that heterogeneity between cells in WT sperm is due to active demethylation by DME and ROS1 (Fig. 2C). Together, these data demonstrate that Dynamic GbM loci are actively demethylated in a subset of all cells in somatic, meristem and germline tissues, and consequently are not demethylated in a specific lineage or developmental context.

**Figure 2:**
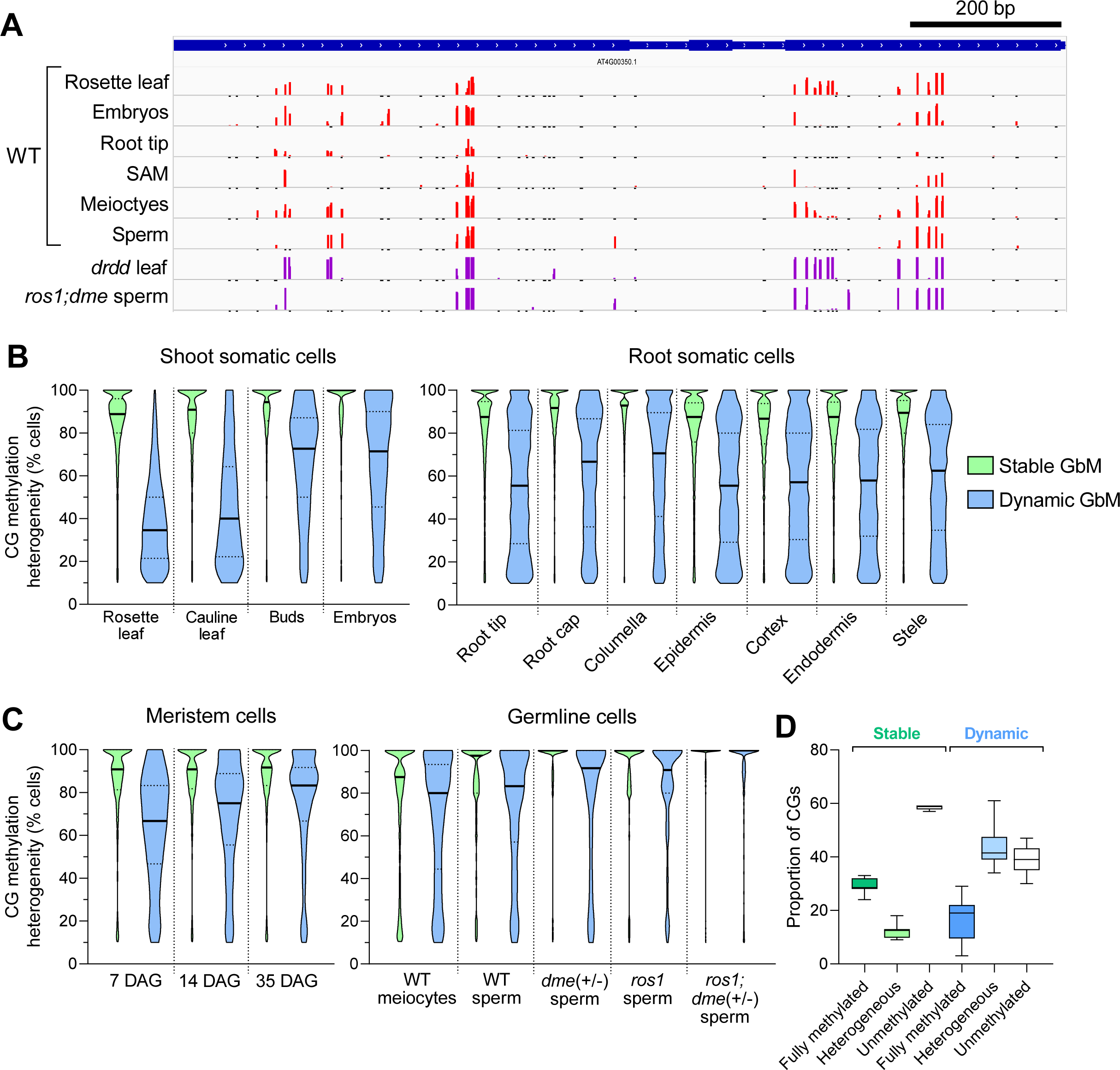
Dynamic GbM genes exhibit methylation heterogeneity in all cell and tissue types. A) Representative genome browser snapshot showing heterogeneously methylated CGs at a Dynamic GbM locus in a variety of cell and tissue types (scale = 0-100%). (B and C) Violin plots showing the distribution of methylation heterogeneity levels of CGs within Stable and Dynamic GbM genes (methylated domains only) from various somatic cell and tissue types (B), and meristem and germ cells (C). Solid lines represent the median and dashed lines represent the interquartile range. All Dynamic GbM violins are significantly different to Stable GbM violins (p=<0.0001), with the exception of *ros1; dme* (+/−) sperm cells. D) Proportion of fully methylated, heterogeneously methylated and unmethylated CGs across all tissue and cell types for both Stable and Dynamic GbM gene bodies.

### De novo methylation is required to maintain dynamic GbM

As Dynamic GbM genes are demethylated in the germline, we reasoned that they must also be subject to *de novo* methylation to counteract DRDD activity. If Dynamic GbM genes are not also targeted by one or more methyltransferases, then DRDD activity would ultimately remove all methylation, and the methylation patterns of dynamic GbM loci would not be stable over inheritance. We observe Dynamic GbM in WT samples from multiple independent laboratories [10,23,33], as well as distinct ecotypes [34] (Figure S1), so the methylation patterns of Dynamic GbM loci must therefore be robust against many generations of inbreeding. To identify the methyltransferase pathway that counteracts DRDD to maintain Dynamic GbM, we examined Dynamic GbM methylation patterns in mutants of the major de novo methyltransferase pathways, RNA-directed DNA methylation (RdDM) [33] and Chromomethylases 2 and 3 (CMT2/CMT3) [33,35]. Unexpectedly, we observed no evidence that these de novo pathways act at Dynamic GbM loci, with quadruple *ddcc* mutants exhibiting near-identical methylation levels and cellular heterogeneity to wild-type (Fig. 3A-B). *De novo* methylation of Dynamic GbM loci must therefore occur by an unknown mechanism. We therefore hypothesized that the maintenance methyltransferase MET1 may exhibit *de novo* activity at these loci. *De novo* methylation by MET1/DNMT1 is not unprecedented – it has been observed in mammalian cells [36], and *de novo* CG methylation has been observed at an introduced transgene within Arabidopsis [37]. Recently, *de novo* CG methylation was also reported in inbred *ddcc* lines [38]. Heterozygous *met1* mutants lose the vast majority of gene body methylation [33], including at most Dynamic GbM genes (Fig. 3C). We therefore chose to exploit this system to perform a genetic experiment to test whether reintroduction of homozygous WT MET1 alleles is sufficient to incur *de novo* methylation at Dynamic GbM loci.

**Figure 3:**
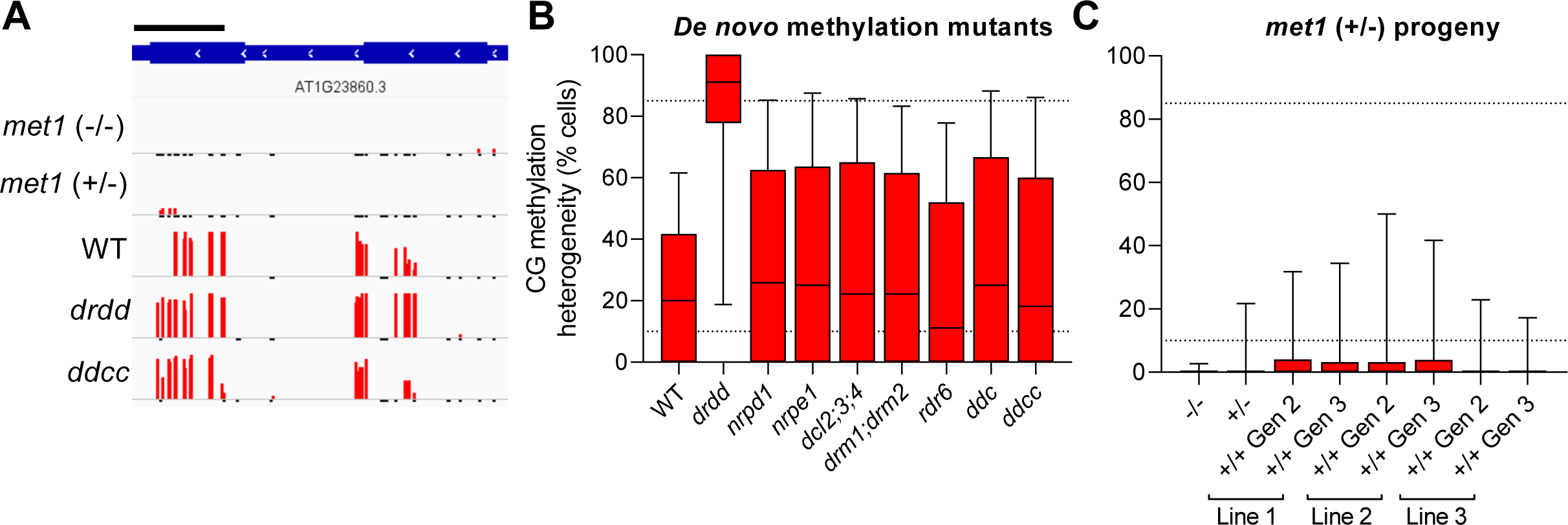
The Dynamic GbM state is maintained by MET1. A) Representative genome browser snapshot of a Dynamic GbM locus in *met1*, *ddcc* and *drdd* mutants. Scale bar = 200 bp. B) Boxplots showing the cellular heterogeneity of CGs within Dynamic GbM genes in WT, *drdd* mutants and multiple de novo methylation mutants. C) Boxplots showing the cellular heterogeneity of CGs within Dynamic GbM genes in homozygous (-/-) and heterozygous (+/−) *met1* mutants, as well as multiple WT-segregant progeny from a selfed *met1* heterozygote. In B and C, horizontal lines = median, box = interquartile range and whiskers = 95^th^ percentile. Horizontal dashed lines represent the cutoffs used to identify heterogeneously methylated CGs (10-85% methylation across all cells).

Heterozygous *met1* mutants were self-fertilized, and two subsequent generations of WT-segregant (+/+) progeny were selected for whole genome methylation sequencing by enzymatic methyl-seq (EM-seq). As has been previously reported, the memory of DNA methylation patterns maintained by MET1 is initially lost in WT progeny of heterozygous mutants [33]. Similarly, we did not see complete reintroduction of Dynamic GbM in WT segregant plants (Fig. 3C). While we did observe a modest gain in methylation at a subset of CGs within Dynamic GbM genes in two out of three wild-type segregant lines, this enrichment was not statistically significant (Fig. 3C). Furthermore, inbreeding WT segregant lines for an additional generation did not result in further restoration of methylation at Dynamic GbM loci (Fig. 3C). We then further assessed the number of putative de novo methylation events per CG across all cells (e.g. the presence of one or more methylated reads per CGs) at Dynamic GbM and Stable GbM gene bodies. As expected, CG methylation events were very rare at both types of GbM genes in homozygous *met1* mutants (Additional file 2: Fig. S3). In *met1* heterozygotes and WT-segregants, we observed that de novo methylation was approximately 25% more common at Dynamic GbM genes that Stable GbM genes (Additional file 2: Fig. S3). We also observed rare but heritable CG methylation events at individual GbM genes, which varied from individual CGs to near-WT restoration of methylation patterns (Additional file 2: Fig. S4). Together, these results suggest that de novo methylation by MET1 at an unmethylated locus is rare. If MET1 does maintain the Dynamic GbM state through *de novo* methylation activity, then it likely occurs when MET1 is recruited to the replication fork by the presence of hemimethylated CGs, or adjacently methylated CGs. Our data do not conclusively prove that MET1 maintains the Dynamic GbM state and thus leaves open the possibility that unknown mechanisms could operate at these loci.

### Dynamic GbM loci are conserved across distant lineages

As Dynamic GbM loci are clearly maintained over many generations of inheritance and between divergent ecotypes of *Arabidopsis thaliana* (Additional file 2: Fig. S2), we sought to determine whether the Dynamic methylation state was conserved over evolutionary time. Evolutionary conservation of the Dynamic GbM state in distant lineages would suggest that the targeting of methylation/demethylation pathways to these loci in all cells serves an adaptive benefit. We isolated the methylation profiles of the most likely homologous genes from five additional genomes representing lineages that diverged from *Arabidopsis thaliana* between 6 and 117 million years ago (Additional File 3) [39,40]. Homologs were identified by blast homology and further filtered using two criteria: 1) only homologs that were classified as one-to-one orthologs by Inparanoid orthology analysis [41] were included, to avoid misidentification of homologs in gene families with duplications. 2) homologs possessing >1% non-CG methylation across the gene body were excluded, in order to remove genes targeted by de novo methylation pathways, which could impact CG methylation heterogeneity. Even in species that diverged >100 million years ago, intermediate methylation levels at individual CGs (representing cellular heterogeneity in methylation) were clearly observable at orthologous genes to the Dynamic GbM loci observed in Arabidopsis (Fig. 4A). To verify that the frequency of intermediately methylated CGs at Dynamic GbM homologs (likely orthologs) were enriched relative to the number expected by chance, we compared homologs to the set of 581 Dynamic GbM genes against homologs to ten randomly selected sets of 581 genes. Both the number and proportion of heterogeneously methylated CGs within Dynamic GbM homolog gene bodies were substantially higher than the number and proportion observed across the gene bodies of randomly selected genes (Fig. 4B). For all species analyzed, the number of heterogeneously methylated CGs was >2-fold higher than in randomly selected genes. These data strongly suggest that the loci targeted by DRRD and MET1 are similar across distant evolutionary lineages, and that the Dynamic GbM epigenetic state may offer an adaptive benefit to certain loci.

**Figure 4:**
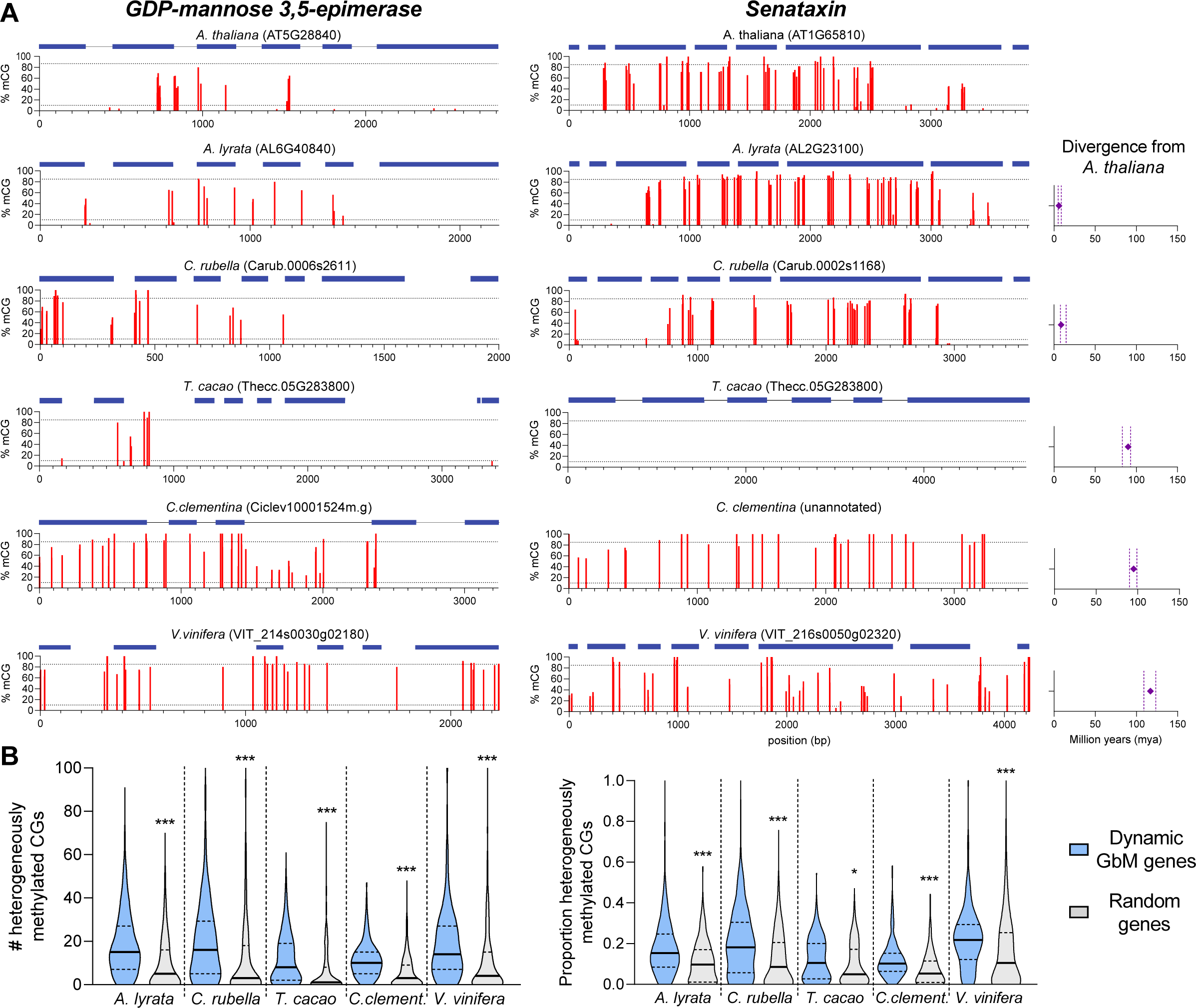
Evolutionary conservation of the Dynamic GbM state. A) Representative Dynamic GbM gene homologs from six different species. Each red bar represents a single CG, and the bar height is the percent methylation heterogeneity across cells. Dashed lines represent the cutoffs used to call heterogeneously methylated cytosines (10-85% of cells). Rightmost panel: the estimated divergence time between species and *Arabidopsis thaliana*, obtained from TimeTree [70]. Points represent the median divergence time estimate, and vertical dashed lines represent the 95% confidence interval. B) Violin plots showing the number (left panel) and proportion (right panel) of heterogeneously methylated CGs across all homologs of Dynamic GbM genes, and ten identically-sized groups of randomly selected genes. Solid lines represent the median and dashed lines represent the interquartile range. *p = <0.05, ***p = <0.0005 (two-tailed t-test).

### Dynamic GbM is associated with a regulatory chromatin state and enhanced gene expression plasticity

As the Dynamic GbM state is conserved over evolutionary time, we sought to understand the biological purpose of this pattern of epigenetic modification. Consistent with our previous study [10], we observed a subset of Dynamic GbM genes are differentially expressed between WT and *drdd* (Additional file 2: Fig. S5). However, these expression changes do not reflect a broad pattern of expression change across all Dynamic GbM genes, leading us to conclude that Dynamic GbM’s primary role is not to consistently alter expression levels, similar to the consensus view on Stable GbM [9,27,28], we did not identify any strong associations between Dynamic GbM and broad patterns of gene expression. Although a small number of Dynamic GbM genes exhibited altered transcript abundance in *drdd* and *met1*+/− mutants (28 genes in *drdd* & 37 non-overlapping genes in *met1*+/−), overall transcriptional output (in standard growth conditions) was largely unaltered in the absence of Dynamic GbM (Additional file 2: Fig. S5). We also found that Dynamic GbM is associated with genes that exhibit a range of expression levels, whereas Stable GbM is predominantly associated with moderate-highly expressed loci (Additional file 2: Fig. S5).

To better understand the possible function of Dynamic GbM beyond impacting gene expression levels, we examined the chromatin state of individual exons exhibiting Dynamic GbM, compared with exons exhibiting a Stable GbM state and genomic total exons as a control. Coding regions are typically associated with the histone marks H3K4me1 and H3K36me3, which consequently correlate with high expression levels, elevated DNA repair, and low mutation rates [42,43]. Both of these histone marks were enriched in total exons relative to the genomic average, and enriched further still in Stable GbM exons (Fig. 5A), consistent with their proposed function as stably expressed, functionally important (e.g. housekeeping) constitutive genes [25,27]. In contrast, Dynamic GbM exons exhibited reduced presence of these typical coding region histone marks, instead showing a significant enrichment in H3K4me2, as well as increased levels of H3K4me3 and histone acetylation marks, such as H3K9ac, H3K23ac, H3K27ac and H3K56ac, all of which are typically associated with 5’ regulatory regions. Exons are typically depleted for the histone mark H3K4me2 which is associated with a regulatory promoter chromatin state [44]. Consequently, we observed a much greater degree of overlap between Dynamic GbM genes and intergenic chromatin states (Fig. 5B-C) – including those normally associated with distal promoter sequences(chromatin state 2 as classified by Sequeira-Mendes et al [44]) – compared to Stable GbM genes. The majority of genes in the genome, including Stable GbM genes, display a single peak of H3K4me2 at the promoter and transcriptional start site, which is replaced by H3K4me1 within the coding region. In Dynamic GbM genes, we observed elongated H3K4me2 domains, which encompassed the entire gene body, or occasionally additional distinct H3K4me2 peaks that coincided with the cytosines displaying dynamic methylation turnover (Fig. 5F, Additional file 2: Fig. S6). While Dynamic GbM genes are typically shorter than Stable GbM genes (Fig. 5D), the enrichment of H3K4me2 is independent of gene length (Fig. 5E). A similar association was also observed for the histone acetylation marks H3K9ac, H3K23ac, H3K27ac and H3K56ac, which are also typically associated with intergenic regulatory chromatin. While these histone acetylation marks are typically localized to the 5’ end of genes, we observed a significant enrichment in Dynamic GbM genes compared to Stable GbM genes of the same length (Additional file 2: Fig. S6). Together, these data suggest that Dynamic GbM genes possess an unusually high enrichment of regulatory “promoter-like” chromatin within the gene body.

**Figure 5:**
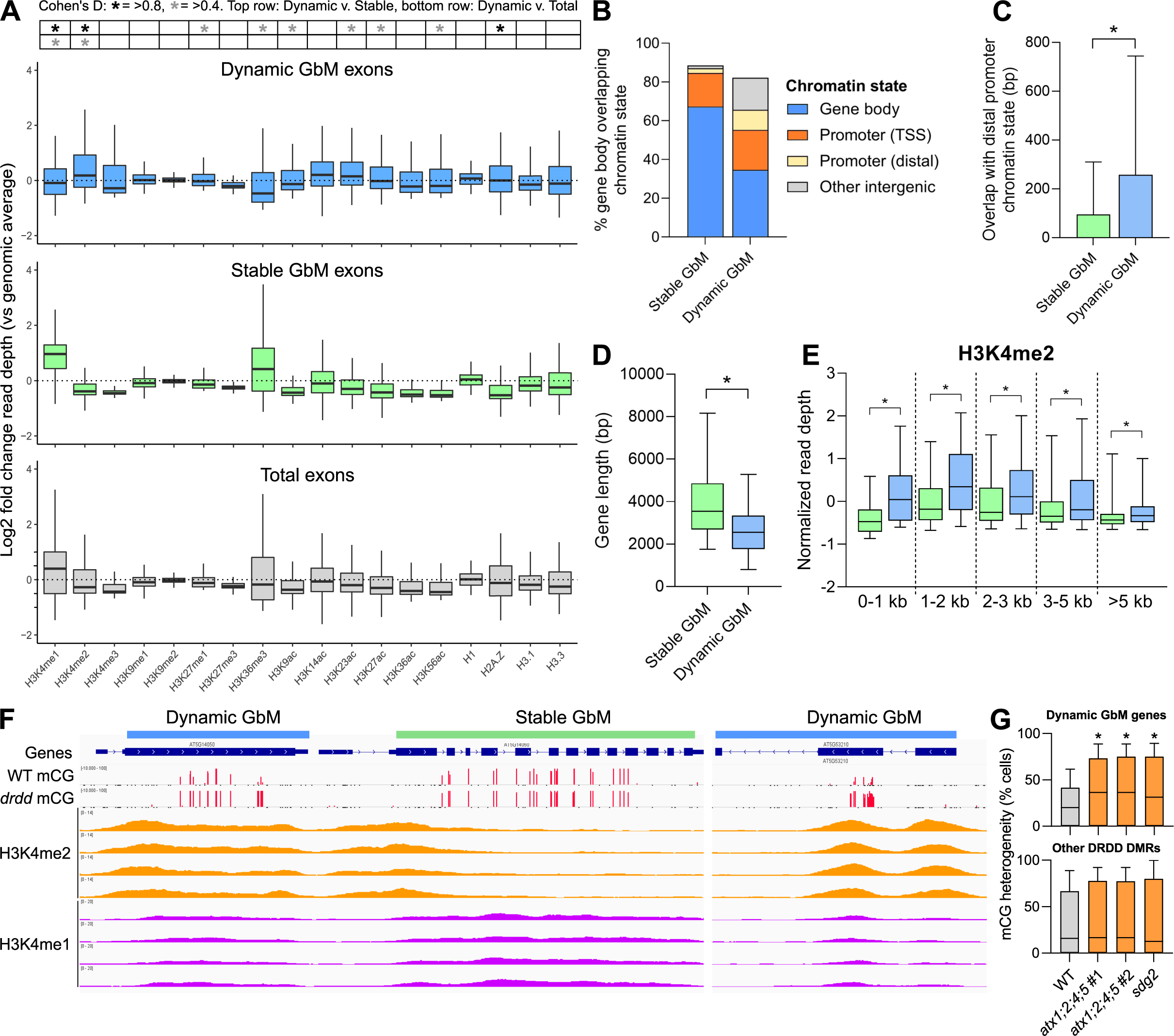
The association between Dynamic GbM and regulatory chromatin marks. A) Boxplots showing the normalized read depth of ChIP-seq data of fourteen histone modifications (data from [42]) within exons overlapping Dynamic GbM domains (blue), Stable GbM domains (green) or total exons (grey). Horizontal lines = median, box = interquartile range, whiskers = 5^th^ and 95^th^ percentiles. The red box highlights model gene body (H3K4me1) and promoter (H3K4me2) histone marks. Statistical test for effect size = Cohen’s D. B) The percentage overlap between Stable and Dynamic GbM gene bodies and four chromatin states identified by [44]. C) The total base pair overlap between Stable and Dynamic GbM gene bodies and a chromatin state typically enriched in distal promoters. D) Boxplot showing the distribution of gene body lengths of both Stable and Dynamic GbM genes. E) Boxplot showing the normalized read depth of H3K4me2 ChIP-seq data at Stable and Dynamic GbM genes separated into bins based on gene body length. For C,D,E: *p = <0.0001 (two-tailed parametric t-test). F) Genome browser snapshot of raw H3K4me1 and H3K4me2 ChIP-seq data at Stable and Dynamic GbM genes. Tracks represent WT data from four independent studies. Data obtained from [69]. G) Boxplots showing the cellular heterogeneity of CGs within the methylated domains of Dynamic GbM genes in WT and two mutant backgrounds with reduced H3K4me2, *atx1;2;4;5* and *sdg2*. *p = <0.0001 (non-parametric two-tailed t-test).

To further explore if a functional relationship exists between the presence of regulatory chromatin and Dynamic GbM, we re-analyzed publicly available DNA methylation data from histone methyltransferase mutants *sdg2* [33], and quadruple *atx1;2;4;5* mutants [45], which are both associated with the methylation of H3K4 and exhibit reduced H3K4me2 and H3K4me3 genome-wide. In both *sdg2* and *atx1;2;4;5* mutants, DNA methylation was increased at Dynamic GbM genes (Fig. 5G), suggesting that the regulatory chromatin state may play a role in DRDD targeting. Methylation increases were not observed at other known target loci of DRDD, suggesting this interaction is specific to Dynamic GbM, and not due to altered regulation of DRDD in *sdg2* or *atx1;2;4;5* mutants. Together, these data show that Dynamic GbM is associated with a regulatory, or “promoter” chromatin state, suggesting a novel link between gene body methylation and transcriptional regulation. To our surprise, only a small subset of Dynamic GbM genes were enriched with H2A.Z compared to the genomic average, which has previously been proposed to associated with the gene bodies of highly responsive genes [46]. Only 130 out of 581 Dynamic GbM genes overlapped with 4080 genes defined as H2A.Z enriched [46]. By contrast, the gene bodies of Stable GbM genes were strongly depleted for H2A.Z compared to the genomic average.

Due to the presence of regulatory histone marks within the gene bodies of Dynamic GbM genes, we sought to understand the relationship between the Dynamic GbM state and transcription. Dynamic GbM genes do not show a category-wide change in transcriptional output between WT and mutants with defective MET1 maintenance of GbM or DRDD demethylation of GbM (Additional file 2: Fig. S5). We therefore hypothesized that the Dynamic GbM state does not directly modulate transcriptional output (like RNA-directed DNA methylation or other silencing mechanisms) but rather facilities a more flexible or open-ended regulatory structure, due to the presence of regulatory histone marks. To test this hypothesis, we leveraged large genomics datasets that compare gene expression across a wide range of tissues and cell types [47–49], biotic and abiotic stresses [50]. Using these datasets, we defined gene expression variability using two statistical measures, the coefficient of variation, and the Fano factor (also known as the index of dispersion). Both of these statistical measures are commonly used to measure gene expression dynamic range. The coefficient of variation has been argued to over-estimate variability of lowly expressed genes, whereas the Fano factor may over-estimate variability of highly expressed genes. We therefore considered that a robust relationship between GbM states and expression plasticity should be statistically significant using both measures. Indeed, Dynamic GbM genes exhibited elevated gene expression plasticity compared to both Stable GbM and total genes across 153 tissue/cell type microarray experiments [48] (Figure 6A), 54 tissue-type samples from a single RNA-seq study [49] (Figure S7) and 165 biotic/abiotic stress condition experiments (Figure 6B). Stable GbM genes exhibited low variability across development and variable physiological conditions, consistent with the observed correlative association between stable gene body methylation and consistently expressed housekeeping genes [25,27] (Fig. 6A). Importantly, we observed no overlap between Dynamic GbM genes and genes with a DRDD-target differentially methylated region in their upstream or downstream intergenic regions, and we observed no increase in the gene expression variance of genes associated with intergenic DRDD targets across development (Additional file 2: Fig. S8). This suggests that increased gene expression plasticity is unique to Dynamic GbM genes, not a consequence of DRDD targeting intergenic regions (genes associated with intergenic DMRs did show increased plasticity in stress condition experiments, which is consistent with the known enrichment of proximal transposons -- which are frequent DRDD targets -- in stress responsive genes [51]) We therefore propose that the abundance of regulatory histone modifications within Dynamic GbM genes enables a greater range of possible expression states, in stark contrast to stable methylation, which promotes canalization towards a single, robust and consistent expression state (Fig. 6C). The significantly enriched GO terms for each subtype of GbM are consistent with this. Enriched GO terms for Stable GbM genes include a number of crucial cellular housekeeping functions, such as DNA repair, RNA splicing, nuclei mRNA export and nuclear protein import (Additional file 2: Fig. S9). Conversely, enriched GO terms for Dynamic GbM include functional categorizations consistent with high expression plasticity and dynamic range over development and in response to the environment, such as photosynthesis, response to temperature, and response to stress (Additional file 2: Fig. S9).

**Figure 6:**
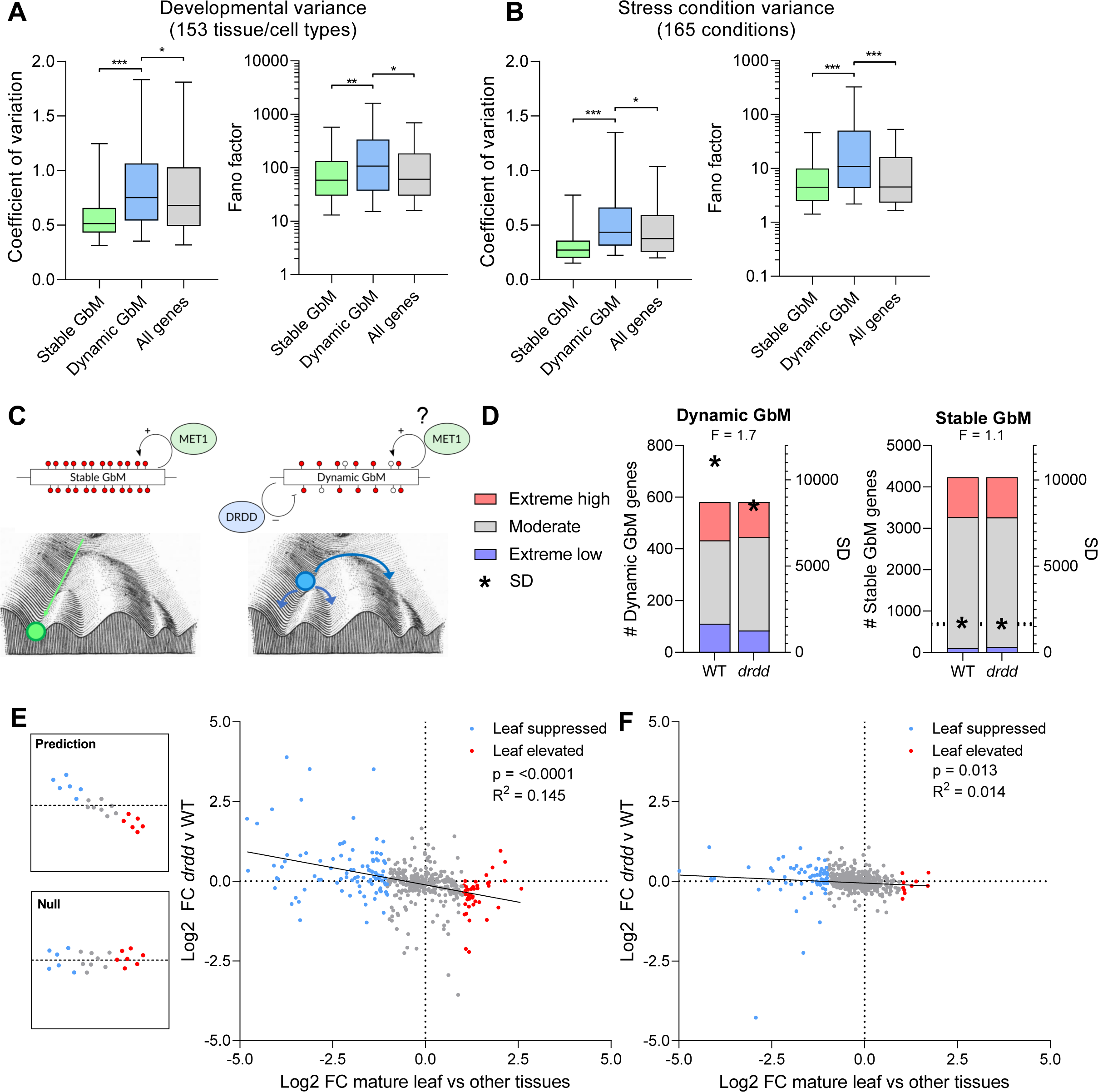
Dynamic GbM is associated with increased gene expression plasticity. A) Boxplots showing the coefficient of variation and Fano factor of expression levels for all genes, Stable GbM, and Dynamic GbM genes across development and B) diverse physiological stress conditions. Horizontal lines = median, box = interquartile range, whiskers = 5^th^ and 95^th^ percentile. ***p=<0.0005, **p=<0.005, *p=<0.05. C, two-tailed t-test) Schematic showing the proposed impact of Stable and Dynamic GbM on gene expression plasticity, represented as Waddington’s landscape. Stable GbM genes exhibit low variance, suggesting canalization into a single, robust gene expression state. Dynamic GbM genes exhibit high variance, suggesting the capacity to access multiple possible gene expression states. D) Proportion of Dynamic and Stable GbM genes in high and low expression deciles in WT and *drdd* mutants. Standard deviation of Dynamic and Stable genes is represented by asterisks. F denotes an F-test for equality of variance (Dynamic GbM p = <0.0001, Stable GbM p = 0.1). E) Scatterplot showing log2 fold change in expression of Dynamic GbM genes between leaves vs other tissues in WT (x-axis) compared to log2 fold change between WT and *drdd* leaves. Loss of gene expression plasticity is predicted to confer a significant inverse relationship. F) Scatterplot comparing equivalent fold-change values for a control set of Stable GbM genes. R^2^, p-value and line of best fit from linear regression model.

One prediction of this model is that disruption of methylation dynamics at dynamic GbM loci should impact gene expression plasticity. *drdd* mutants possess consistently high methylation levels at Dynamic GbM genes in the majority of all cells, similar to Stable GbM genes in WT (Fig. 1E). We therefore sought to test if *drdd* mutants exhibit reduced variance in Dynamic GbM expression levels compared to WT. Consistent with our model, Dynamic GbM genes exhibited a reduced dynamic range in *drdd* mutants, with fewer genes in the highest and lowest expression deciles and aa 30% lower standard deviation to WT (Figure 6D), whereas the standard deviation in Stable GbM gene expression was unchanged. To investigate this further, we sought to determine if the bi-directional differences between Dynamic GbM gene expression in leaf tissues WT and *drdd* (Additional file 2: Fig. S4) could be explained by reduced gene expression plasticity in *drdd* mutants. If DRDD activity is functionally required to enable gene expression plasticity during WT development, then Dynamic GbM genes highly expressed in leaves should exhibit reduced expression (closer to the mean for all tissues) in *drdd*, whereas lowly expressed genes should exhibit higher expression. We therefore compared the Log2 fold-change of Dynamic GbM genes in rosette leaves compared to all other tissues using a 54-tissue study of the Arabidopsis transcriptome [49], and compared this against the Log2 fold-change in expression between WT and *drdd* leaves (Fig. 6E). This revealed a clear and highly significant (linear regression) inverse relationship between the expression change of Dynamic GbM genes in *drdd* and their natural expression state in WT leaves (Pearson’s correlation coefficient = −0.4, linear regression R^2^ = 0.145). For example, genes with peak expression in WT leaves vs other tissues show reduced expression in *drdd* leaves, consistent with reduced gene expression plasticity. While an R^2^ value of 0.145 might seem small, we note that *drdd* is a highly pleiotropic mutant with many phenotypic and transcriptomic changes. Explaining 14.5% of the variance in the transcriptome profile of *drdd* Dynamic GbM genes solely from the expression enrichment of WT leaves vs other tissues is a notable and highly significant (p = <0.0001) result. Conversely, a control group of Stable GbM genes exhibit almost no relationship between the two (Fig. 6F). This finding further supports our model – that the turnover of DNA methylation patterns at Dynamic GbM loci is associated with enhanced gene expression plasticity, and also strongly suggests a functional role for demethylation by DRDD in enhancing plasticity of expression states.

## DISCUSSION

DNA methylation within the bodies of genes has been a well-known feature of eukaryotic genomes for several decades [26,27,52]. Despite this, the precise function of gene body methylation has remained unclear. A positive correlation between gene body methylation and robustly expressed genes has been well established in both animals and plants [27,52], with genes that occupy the extremes of the distribution of expression states typically being unmethylated in their gene body [27]. In this study, we have presented further evidence that gene body methylation functions by modulating the possible positions within “expression space” that genes can occupy. Stable gene body methylated genes (in which CGs are methylated in every cell) exhibit a dramatically lower gene expression plasticity across development and diverse physiological conditions than the genomic average (Fig. 6). A recent high-throughput genetic knockout study also supports this model. Genetic knockout of the CG methylation maintenance methyltransferase MET1 in 18 different wild-type Arabidopsis accessions resulted in a dramatic increase in overall expression variability between genotypes, suggesting that CG methylation functions in canalizing transcriptional activity, or reducing variability [53]. Interestingly, studies of the responses of both animals and plants have also identified GbM as a correlate of gene expression plasticity. A study of two seagrass species showed that the genes that display low expression plasticity in response to environmental stress share properties of Stable GbM genes and are predicted to be methylated [54]. A similar association has been observed in corals transplanted between high and low fitness environments, where GbM is proposed to manage the balance between low/un-methylated environmentally-responsive genes and transcriptionally stable GbM genes [55]. Lastly, intergenic DNA methylation has been shown to prevent spurious transcription initiation in mammals [56], which may symptomatic of an epigenetic state that promotes robust pol II elongation and reduced transcriptional noise. Understanding the molecular mechanism through which Stable GbM reduces expression variability will be an interesting avenue for future research.

Our study identifies a new sub-type of GbM locus within plant genomes, an exception that helps demonstrate the rule associating GbM with reduced gene expression plasticity. This sub-type of GbM, which we term Dynamic GbM, is distinguished by the targeting of methylated CGs by the DRDD DNA demethylation pathway in all cell and tissue types. Dynamic GbM genes display greatly elevated gene expression plasticity in comparison to their Stable GbM counterparts, strongly suggesting that enzymatic modification of the Stable GbM state disrupts its primary function in restricting expression plasticity. One consequence of this DNA demethylation activity is the generation of cellular heterogeneity in DNA methylation patterns and the production of unmethylated CGs within stem and gamete cells. In order to be stable over inheritance and evolution, Dynamic GbM loci must therefore also be re-methylated by one or more methyltransferase enzymes, most likely the maintenance methyltransferase MET1. The dynamic activity of demethylase and methyltransferase enzymes at these loci therefore functions as a constant source of methylation turnover, in stark contrast to the cellular consistency of Stable GbM genes. We therefore propose that dynamic methylation turnover in all cell types is a mechanism that could enable greater expression plasticity over the development of the entire organism. As the targeting and expression of methylation modifier pathways is not cell or tissue-type specific in plants, we reason that they target Dynamic GbM genes universally, so that increased gene expression plasticity is available when adaptively beneficial. Consequently, a Dynamic GbM gene may be targeted by methylation writers and erasers even in cell types or conditions in which the gene is not expressed.

While our study offers strong evidence for an association between GbM and gene expression plasticity, this proposed function is largely based on a correlative association, similar to prior hypotheses of GbM function. To move towards a mechanistic understanding of the roles of Dynamic and Stable GbM, we performed a detailed examination of the chromatin states of each. Our results show a striking association between Dynamic GbM and chromatin marks typically associated with regulatory regions, such as H3K4me2. This is consistent with the increased expression plasticity of Dynamic GbM genes, as genic H3K4me2 is likely to increase the extent to which regulators of transcription can target and modify expression of these loci [57,58]. H3K4me2 has also been shown to be associated with genes with high expression variability in Arabidopsis, as well as increased tissue-specificity across development [59], consistent with our proposed model. Intriguingly, gene-body H3K4me2 has also been reported in human cells, and – consistent with our model – is enriched within genes that show high cell-type specificity across development [60], aka elevated plasticity in possible expression states. This raises the possibility that properties of the Dynamic GbM epigenetic state could be conserved across eukaryote kingdoms. In contrast to Dynamic GbM genes, Stable GbM genes show the strongest genic enrichment of histone marks associated with polymerase II elongation and the “housekeeping gene” chromatin state, namely H3K4me1 and H3K36me3 [44]. Defining the precise molecular relationship between gene body methylation, dynamic methylation turnover, expression plasticity and a regulatory histone modification state is a key future extension of this research. Another recent study has also uncovered a subset of GbM genes that deviate from Stable GbM genes with unconventional histone modifications [61]. These GbM genes are dependent on the chromatin modifier DDM1, and are concurrently enriched for H1, H3K27me3 and H2A.Z, as well as the coding region histone marks normally associated with Stable GbM [61]. The GbM subtype identified by this study is distinct from Dynamic or Stable GbM, and raises the possibility that there are additional sub-types of gene-body methylated epigenetic states to be characterized. Two findings within our study point towards a causative role for DNA methylation dynamics in impacting gene expression variability. First, *drdd* mutants, which cannot generate methylation dynamics at GbM genes, exhibit a reduction in the total dynamic range of Dynamic GbM gene expression states (Figure 6D). Second, a significant portion of the gene expression differences between WT and *drdd* mutants can be explained by loss of gene expression plasticity in Dynamic GbM genes (Fig. 6E). As gene expression plasticity is an important determinant of how organisms adapt to stress, variable environments, and climate change [55,62], discerning the precise ways in which epigenetic states impact plasticity is an important future question.

## CONCLUSIONS

In this study, we define a new feature of the epigenetic landscape in plants. A subset of gene bodies within the genome are targeted by DNA methylation writers and erasers in all cells, which create a dynamic DNA methylation turnover. This dynamic DNA methylation state is associated with a “promoter-like” chromatin state, as well as enhanced gene expression plasticity. Our study also demonstrates a clear link between typical gene body methylation (which is stable across all cells) and a canalized expression state with low plasticity. Overall, our study offers a substantial advance towards understanding the functional role of enigmatic gene body methylation in eukaryotes.

## METHODS

### Plant Growth Conditions

All plants were grown on a 1:1 mix of potting compost and vermiculite in a Percival AR100L3 with 16-hour days at 21-22°C and 60% humidity.

### *met1* mutation segregation

Seeds from a single self-fertilized heterozygous *met1-3* [63] plant (obtained from the Arabidopsis Biological Resource Center – stock number CS16394) were grown for 4 weeks in standard growth conditions as outlined above. A single rosette leaf was collected from individual plants and DNA was extracted as described [64]. The genotype of the *MET1* locus was then assayed using the following primers: METNF2, TAGCCAACAAGTTATCGCTTACTC; METNR2, TTCGCAAACCATTCTTCACAGAGC; TL-4, TAATTGCGTCGAATCTCAGCATCG. Plants that were identified as heterozygous for *met1-3* and homozygous for the WT *MET1* allele were self-fertilized and collected to repeat as described for an additional generation.

### DNA extraction and Enzymatic Methyl Sequencing (EM-Seq)

Leaf tissue from 28-day old plants was collected and flash frozen in liquid nitrogen. Two leaves per sample were ground using a Qiagen TissueLyser II and immediately after 200 µl of CTAB was added and heated to 65°C for approximately 20 minutes. This was followed by the addition of 200 µl chloroform, 10 minute centrifugation and ethanol washes to isolate total genomic DNA. The DNA concentration was quantified using a Qubit Flex fluorometer, and a diluted spike-in was added according to NEB recommendations (1 µl diluted pUC19 (0.001 ng) control DNA and 1 µl diluted unmethylated lambda DNA (0.02 ng) in 0.1X TE, pH 8.0 per 10–200 ng plant sample DNA). Genomic DNA samples with added spike-ins were sonicated to an average fragment size of 300bp using a Covaris S220 Focused Ultrasonicator (80s treatment, 140 peak power, 10 duty factor 200 cycles/burst). Subsequently, samples were incubated with an RNAse cocktail and RNAse was removed using a Qiagen spin column. EM-seq libraries were constructed following the NEBNext® Enzymatic Methyl-seq Kit protocol. Library quality was verified with a KAPA Library Quant Kit and fragment analyzer. The samples were pooled to a concentration of 4 ng/µl and 19.2 nM and sequenced using an Illumina NextSeq 2000 (paired-end 150 x 150 bp) 11 million average reads/sample. Further information on each EM-seq library sequenced in this study is available in Supplemental Table 2.

### Whole genome methylation analysis

Mapping of whole genome bisulfite sequencing and EM-seq libraries was performed as described [10]. In brief, reads were processed with TrimGalore 0.6.6 (Babraham Bioinformatics), trimming 8bp from the 5’ end of reads and enforcing a 3’ end quality score of >25%. Reads were mapped to the Araport11 genome using Bismark 0.22.3 [65], allowing for 1 mismatch per read in the seed region and removing PCR duplicates. Methylation values for each cytosine were calculated using the Bismark methylation extractor function. The efficiency of EM-seq conversion was verified by quantifying the percentage of methylation for reads mapped to the chloroplast. EM-seq conversion rates were >99.7% across all samples.

To classify Stable and Dynamic GbM genes, data from WT and *drdd* mutant bisulfite sequencing and EM-seq libraries [10] was used. Analysis was initially performed on a single high-depth EM-seq replicate to avoid inter-individual variation in methylation profiles. Subsequently, the classification of Dynamic GbM genes was also verified by pooling four bisulfite sequencing replicate libraries, showing that Dynamic GbM identification is robust against variation between replicates of the same genotype. Methylation heterogeneity for each CG was calculated by dividing the number of methylated (deduplicated) reads by the number of total (deduplicated) reads. A violin plot for the genomic CG methylation heterogeneity was plotted (R vioplot function) using methylation heterogeneity values for all individual CGs genome-wide. Based on this violin plot, the vast majority of CGs exhibited <10% or >85% methylation heterogeneity, leading to the following classification: CGs exhibiting a methylation percentage <= 10% were classified as unmethylated, CGs >=85% were classified as fully methylated and CGs methylated between 10-85% were classified as heterogeneously methylated, as each deduplicated read derives from an independent cell in the original sample. Dynamic GbM genes were identified based on two cutoffs: 1) the presence of a WT vs *drdd* differentially methylated region (DMR) within the gene body (DMRs were called based on aggregation of differentially methylated CGs within overlapping 200bp windows, as described in our previous work [10,24]). 2) Dynamic GbM genes required at least 5 CG dinucleotides (10 CG cytosines total) that gained >20% methylation in *drdd* mutants. This second step removed genes possessing only a small number of heterogeneously methylated sites in an otherwise unmethylated or fully methylated gene body. Stable GbM genes were identified as described [9]. These two cutoffs resulted in a list of 581 total Dynamic GbM genes.

The evolutionary conservation of Dynamic GbM was performed by identifying most likely homologs to the 581 Arabidopsis genes using BLAST homology from 5 additional species: *Arabidopsis lyrata*, *Capsella rubella*, *Theobroma cacao*, *Citrus clementina* and *Vitis vinifera*. Homologs were further filtered using two criteria: 1) only homologs that were classified as one-to-one orthologs by Inparanoid orthology analysis were included (a list of homologs for each species is included in Additional File 2), 2) homologs possessing >1% non-CG methylation across the gene body were excluded, in order to remove genes targeted by de novo methylation pathways. Additionally, most likely homologs were also identified for ten sets of 581 randomly selected genes, which were then averaged as a control group. The whole genome methylation profiles of each species were downloaded and remapped from other studies [39,40]. For each Dynamic GbM / random homolog, the number of heterogeneously methylated CGs was isolated using bedtools intersect. Both the number and density of heterogeneously methylated CGs was compared relative to average across 10 sets of randomly selected genes. Statistical significance was calculated using an unpaired two-tailed t-test.

### RNA-seq analysis

Prior to mapping, adapters were trimmed using Trim Galore 0.6.6 (Babraham Bioinformatics), trimming 9 bp from the 5’-end of reads, and enforcing a 3’-end quality of >25%. Reads were mapped to the Araport11 genome using STAR 2.7.1a [66], permitting 0.05 mismatches as a fraction of total read length and discarding reads that did not map uniquely. Differentially expressed genes were identified by running htseq-count 0.9.1 and DESeq2 1.40.1 [67], ensuring a minimum of two-fold change in expression and a Benjamini–Hochberg corrected P-value <0.05. GO term analysis was performed using agriGO v2.0 with default parameters [68].

### Chromatin state analysis

Chromatin state classification for the entire Arabidopsis genome was obtained from Sequeira-Mendes et al [44]. The percentage and base-pair overlap between Stable and Dynamic GbM genes and each chromatin state was calculated using Bedtools 2.28.0 intersect. In Figure 5, “Gene body” refers to chromatin states 3 and 7 combined, “Promoter (TSS)” refers to state 1, “Promoter (distal)” refers to state 2, and “Other intergenic” refers to states 4, 5, 8 and 9 combined. Analysis of histone mark enrichment was performed using publicly available ChIP-seq data as previously described [42]. In brief, multiple WT study ChIP-seq datasets were downloaded from the Plant Chromatin State Database (PCSD) [69] as bigwig files, converted into bedfiles displaying raw read depth for each base pair, and normalized by calculating the log2 fold enrichment for each position compared to the genomic average. Fold enrichment was then averaged across the genomic features analyzed (e.g. Dynamic GbM, total exons etc.) using bedtools merge.

### Expression variance analysis

Gene expression counts for Stable GbM genes, Dynamic GbM genes and Total genes were obtained from large publicly available expression datasets, including the the Bio-Analytic Resource for Plant Biology (BAR) Expression Browser platform [48], and a recent study of 54 tissue types [49]. The coefficient of variation and Fano factor for each gene was then calculated using all available data across tissues/cell types or physiological stress conditions. Data from the BAR platform were analyzed using the mean values for technical/biological replicates for each tissue sample/condition. 133 genes that were identified in both the Stable and Dynamic GbM gene groups were excluded to avoid overlap between datasets. Expression variance was also calculated for genes associated with intergenic DRDD-target DMRs, which were defined as any gene possessing a hypermethylated DMR in *drdd* mutants within 2kb upstream or downstream of the coding region. The dynamic range of Dynamic and Stable GbM genes in *drdd* was analyzed calculating the expression decile of each gene in WT and the corresponding expression decile in *drdd*. To test the change in expression in *drdd* relative to WT tissue-specificity, the log2 fold change was calculated between the mature rosette leaf 6 sample sequenced by Mergner et al [49], which is analogous to our *drdd* leaf RNA-seq data, and all other 53 tissue samples sequenced by Mergner et al. These leaf-fold-change values were then plotted against the log2 fold-change between WT and *drdd* rosette leafs as calculated by DE-seq2. Genes with 0 read coverage in either WT or drdd samples were excluded from this analysis. As a control, the analysis was also performed with 600 randomly selected Stable GbM genes.

## DECLARATIONS

### Ethics approval and consent to participate

Not applicable

### Consent for publication

Not applicable

### Availability of data and materials

All high throughput sequencing data generated in this study is available NCBI Gene Expression Omnibus (GEO) [71].

### Authors’ contributions

CJW and KAT performed experiments. CJW, DD, GJM and BPW performed data analysis. BPW conceived of the study and wrote the manuscript.

## Acknowledgements

We are grateful to Annika Quist for her assistance with mapping RNA-seq data. We are thankful to Bob Schmitz, Steve Jacobsen and PRV Satyaki for helpful feedback and discussion.

## ADDITIONAL INFORMATION

### Supplementary Information

**Additional File 1:** A list of Dynamic GbM and Stable GbM genes identified in this study **Additional File 2:** Supplementary Figures and Tables. **Fig. S1**: Expression of *DRDD* genes across tissues and relationship with methylation levels at Dynamic GbM genes. **Fig. S2**: Dynamic GbM methylation levels across 100 Arabidopsis accessions. **Fig. S3**: De novo methylation events in *met1* mutants and WT segregants. **Fig. S4**: Example de novo methylation events in *met1* mutant segregants at GbM loci. **Fig. S5**: RNA-seq analysis of Stable and Dynamic GbM genes. **Fig. S6**: Enrichment of histone acetylation in Dynamic and Stable GbM genes of different lengths. **Fig. S7**: Gene expression plasticity measured from high-throughput RNA-seq study of 54 tissue types. **Fig. S8**: Gene expression plasticity of genes closely linked to intergenic DRDD targets. **Fig. S9**: Enriched GO-terms in Dynamic and Stable GbM genes. **Table S1**: A list of all external datasets analyzed in this study. **Table S2**: Properties of EM-seq libraries sequenced in this study.

