## Additional file 2 for "Dynamic DNA methylation turnover in gene bodies is associated with enhanced gene expression plasticity in plants"

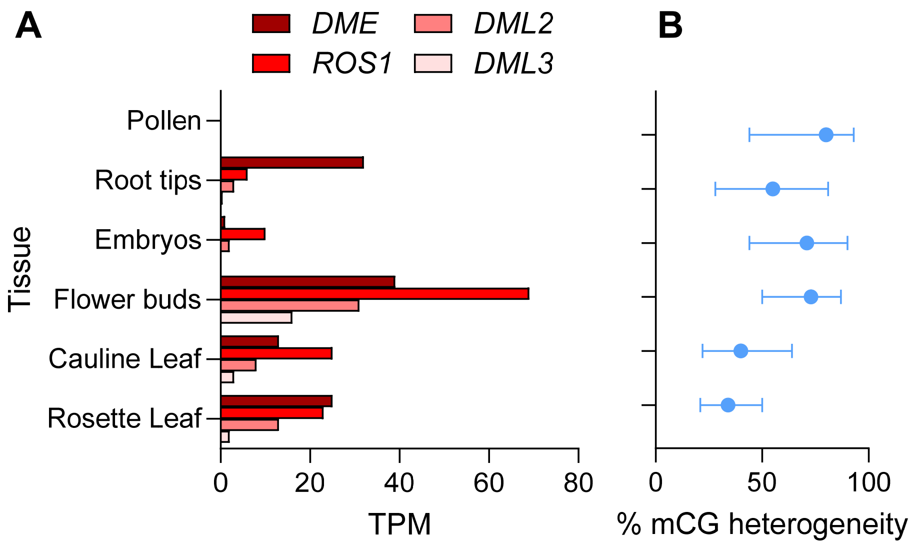


**Figure S1:** Expression of DRDD genes does not correlate with Dynamic GbM methylation heterogeneity. A) Average expression level of *DME, ROS1, DML2* and *DML3* in six tissue types measured by RNA-seq. B) CG methylation heterogeneity of Dynamic GbM genes in six tissue types. Dots represent median values, error bars represent interquartile range.

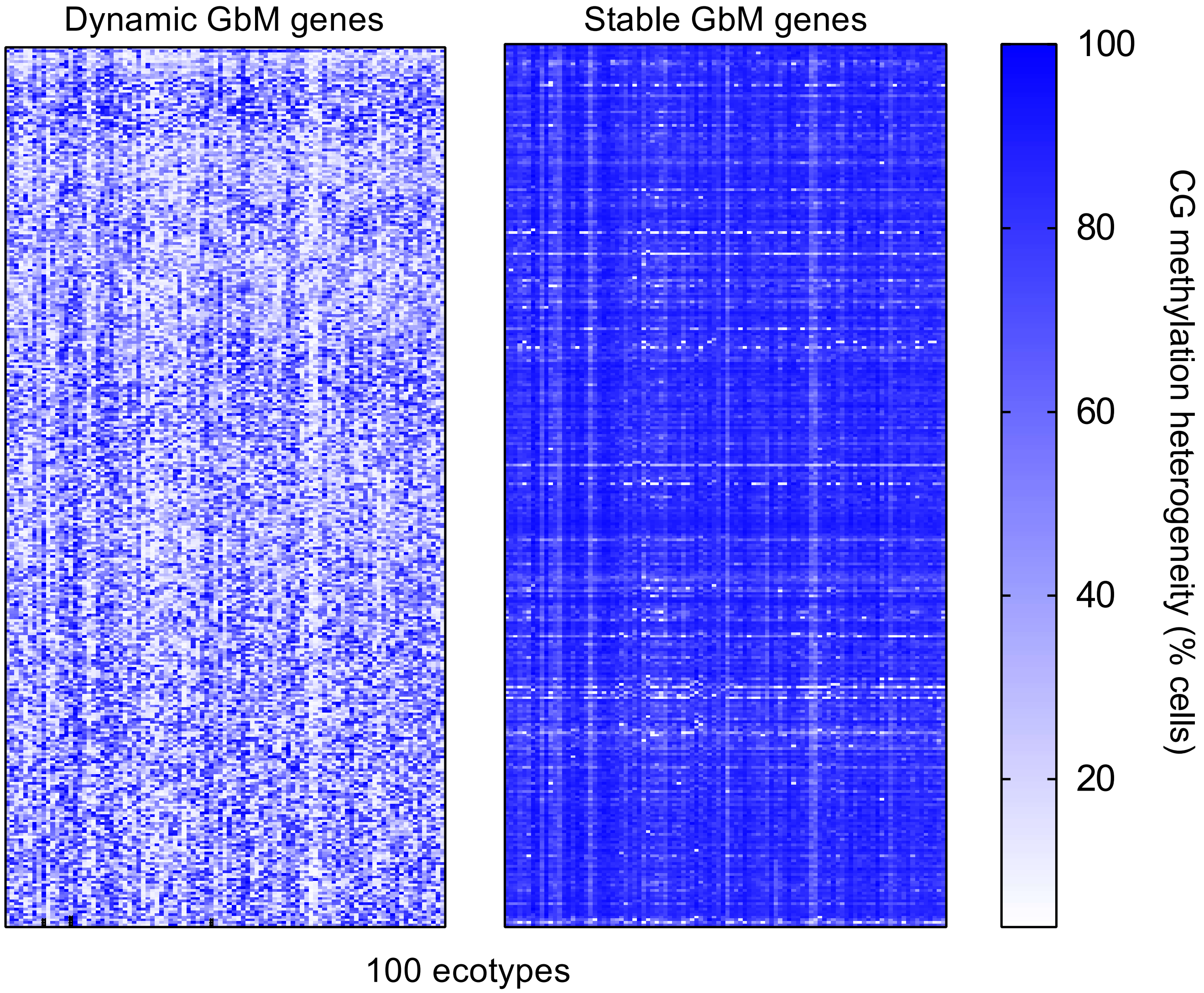


**Figure S2: Dynamic GbM is consistent across Arabidopsis ecotypes**. A heatmap showing CG methylation heterogeneity averaged across the gene bodies of Dynamic GbM and Stable GbM genes in 100 randomly selected ecotypes from the 1001 genomes project [34].

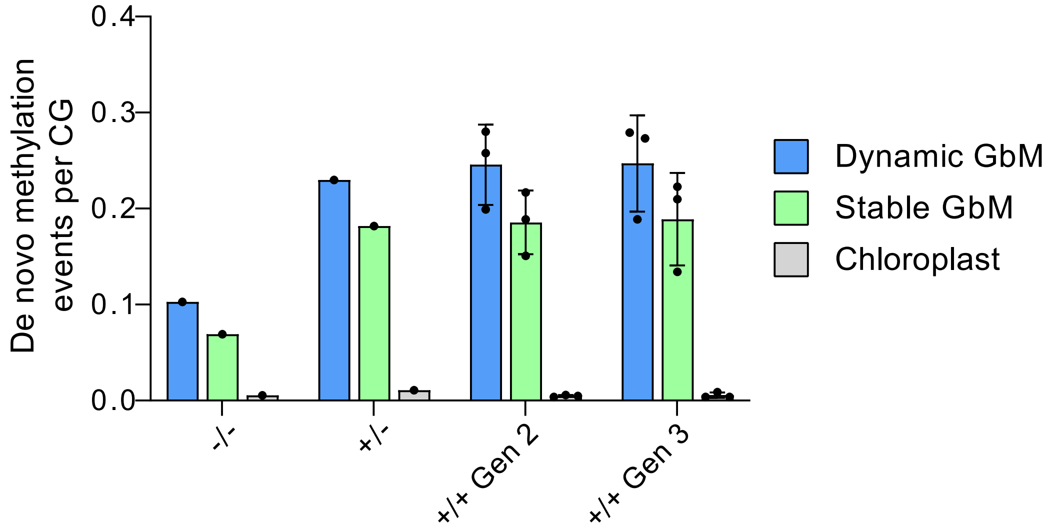


**Figure S3: *De novo* methylation events in *met1* mutants and WT segregants**. The rate of de novo CG methylation events per CG across all cells. Bars represent the mean, error bars represent standard deviation. Individual data points represent independent genetic lines that were sequenced. Chloroplast CG methylation is included a control for the false positive rate from incomplete conversion or sequencing errors.

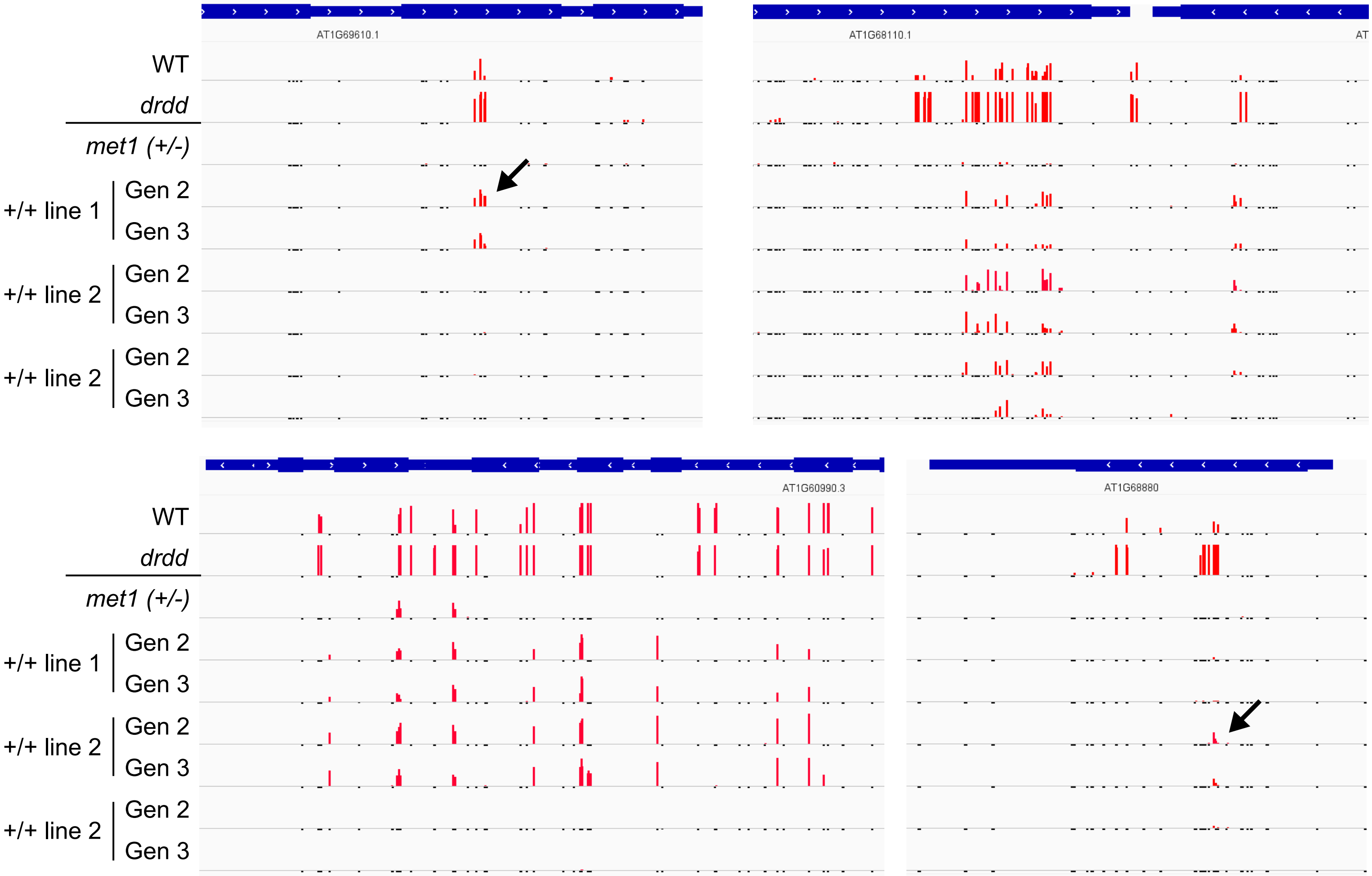


**Figure S4: Example de novo methylation events in *met1* mutant segregants at GbM loci.** Genome browser snapshots of four Dynamic GbM genes in WT, *drdd*, *met1* heterozygotes (+/-) and three independent lines of WT-segregant progeny (+/+).

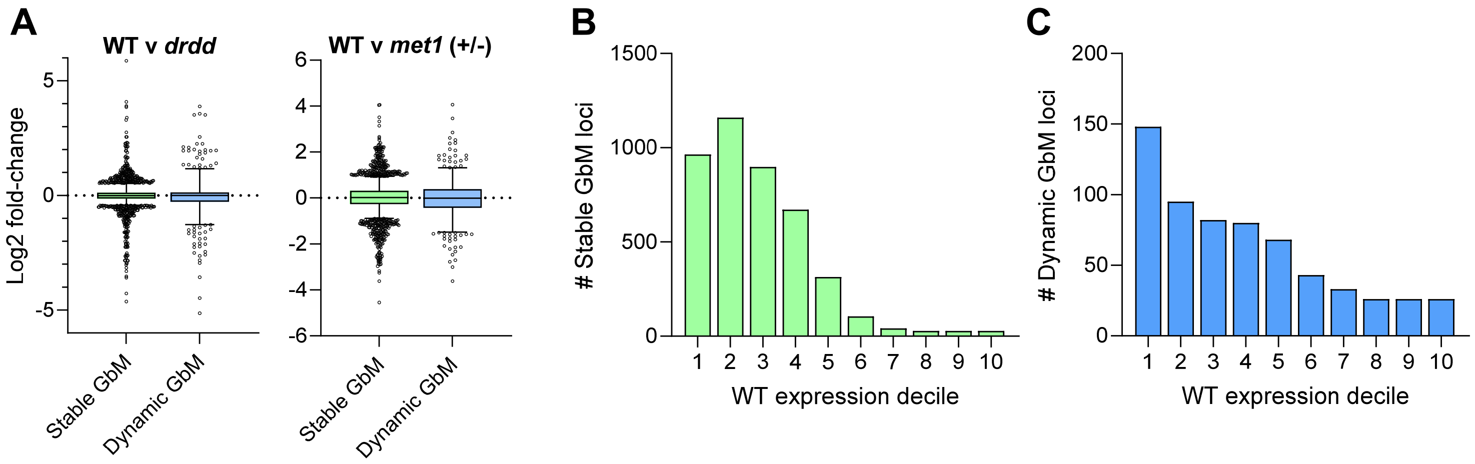


**Figure S5:** RNA-seq analysis of Stable and Dynamic GbM genes. A) Boxplots showing distribution of Log2 fold-change in transcript abundance for Stable and Dynamic GbM genes in WT v *met1* heterozygous mutants (which lose GbM) and WT v *drdd* mutants (which exhibit elevated GbM at Dynamic GbM genes). Horizontal line = median, box = interquartile range, whiskers = 1st and 99th percentile. B) Distribution of Stable GbM genes across WT transcript abundance deciles. C) Distribution of Dynamic GbM genes across WT transcript abundance deciles. In B and C, 1 = highest expression decile, 10 = lowest expression decile.

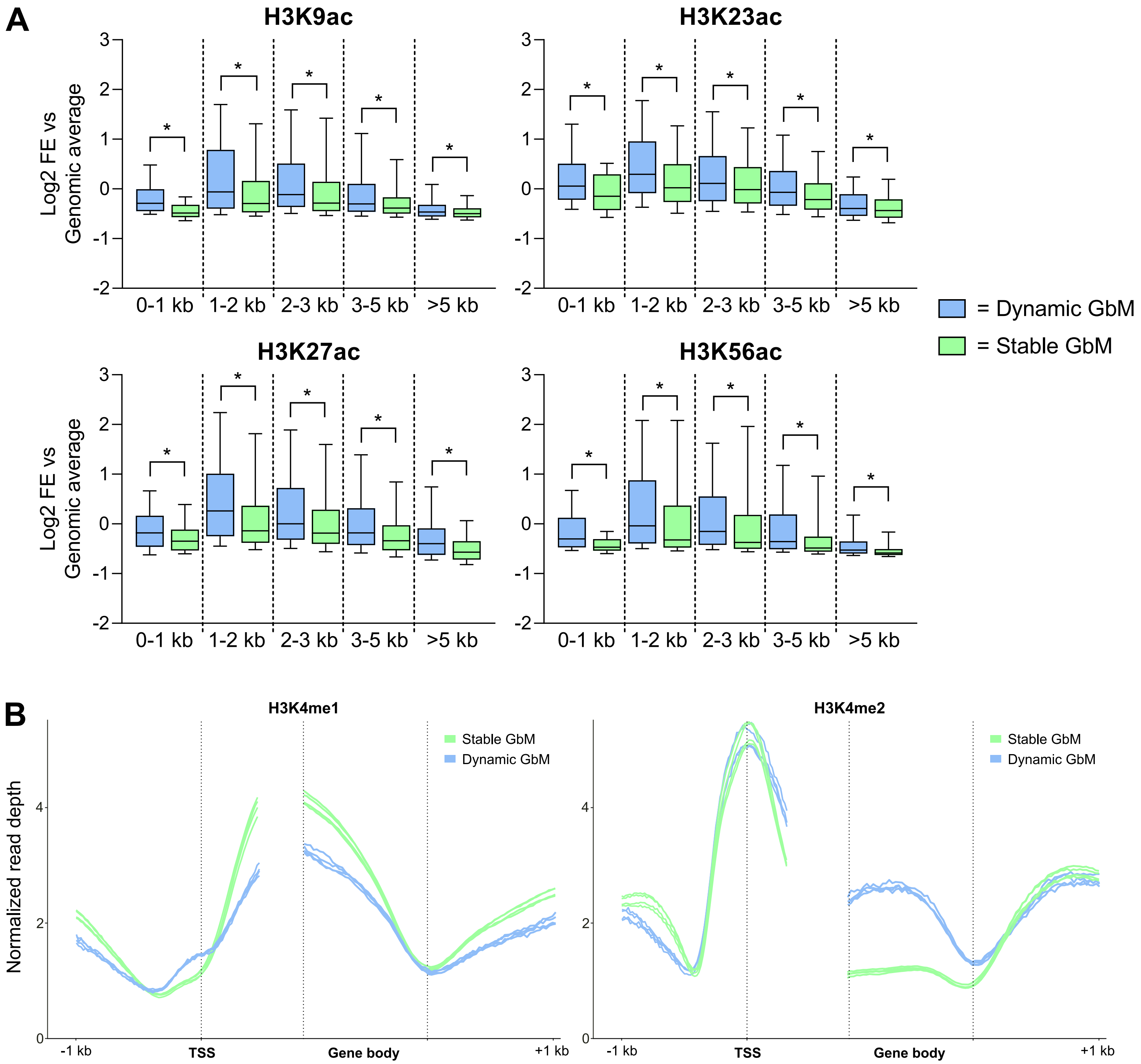


**Figure S6: Enrichment of histone acetylation in Dynamic and Stable GbM genes of different lengths**. A) Boxplots show the log2 fold read depth enrichment of four histone acetylation marks relative to the genomic average (ChIP-seq data was obtained from the plant chromatin state database[69]). Enrichment values are averaged across the whole coding region for each gene. Horizontal line = median, boxes = interquartile range, whiskers = $5^{th}$ & $95^{th}$ percentile. *p = <0.0001 (two-tailed parametric t-test). B) Smoothed moving average plots of ChIP-seq normalized read depth for H3K4me1 and H3K4me2 at Dynamic and Stable GbM genes. Individual lines represent ChIP-seq profiles from indpenendent studies of WT Arabidopsis, obtained from the plant chromatin state database.

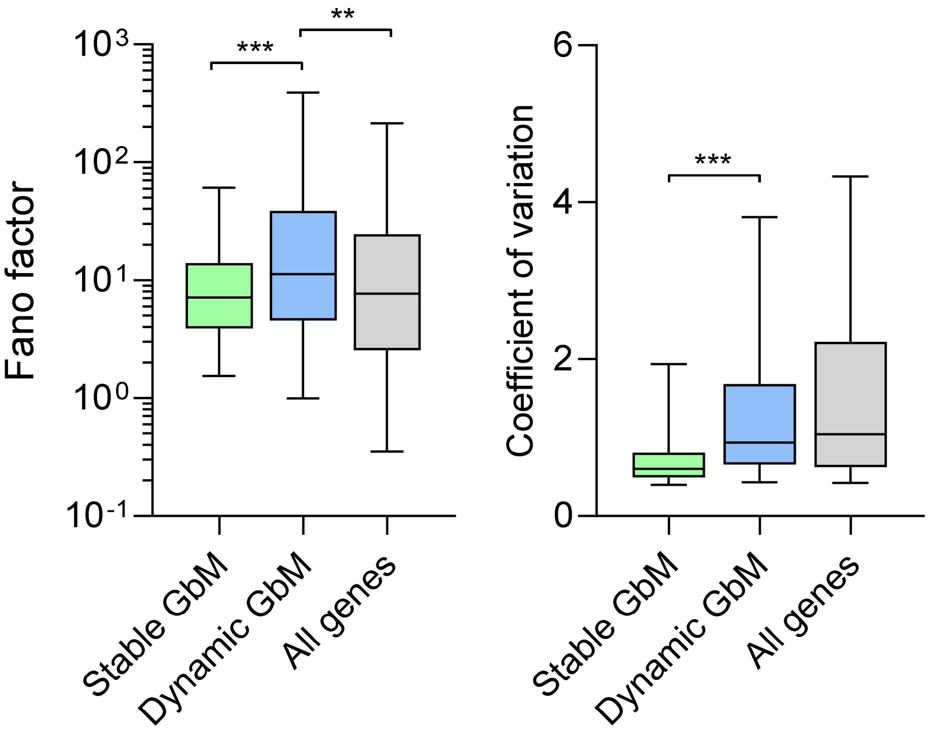


**Figure S7:** Gene expression plasticity measured from high-throughput RNA-seq study of 54 tissue types. Boxplots denote gene expression variability of Dynamic GbM, Stable GbM and total genes measured by the Fano factor (left) and Coefficient of variation (right). Transcript per million values of genes across 54 tissues were obtained from Mergner et al, 2020 [49]. Horizontal lines = median, boxes = interquartile range, whiskers = 5th & 95th percentile. ***p = < 0.0005, **p = < 0.005 (two-tailed parametric t-test).

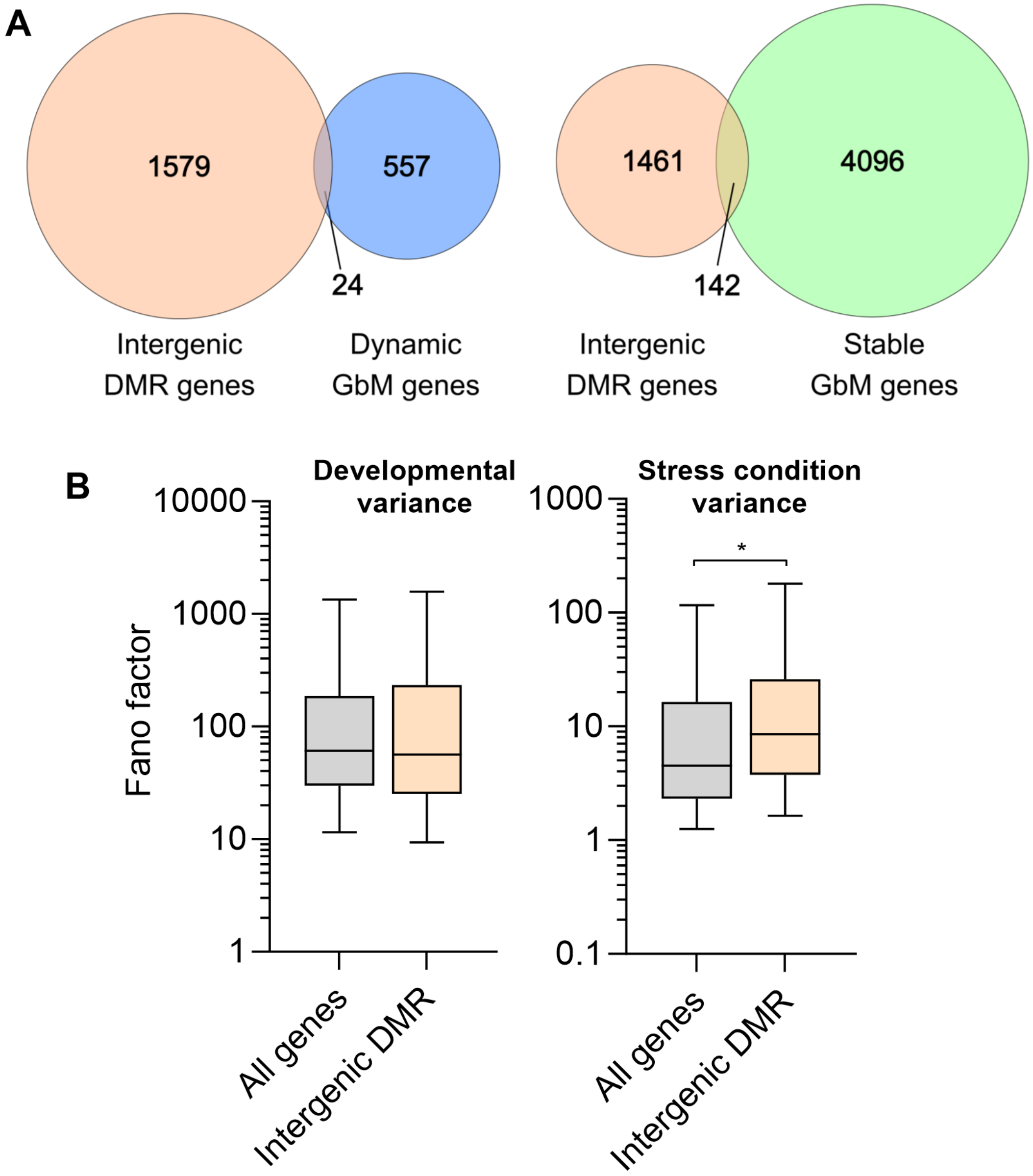


**Figure S8:** Gene expression plasticity of genes closely linked to intergenic DRDD targets.
A) Venn diagrams showing overlap between intergenic DMR genes (defined as a gene harboring a DRDD-target differentially methylated region in intergenic sequences within 2kb upstream or downstream of the coding region) and Dynamic or Stable GbM genes. B) Boxplots displaying gene expression variability as measured by the Fano factor across diverse developmental stages (left) and diverse physiological/stress conditions (right). Horizontal lines = median, boxes = interquartile range, whiskers = 5th & 95th percentile. *p = < 0.05 (two-tailed parametric t-test).

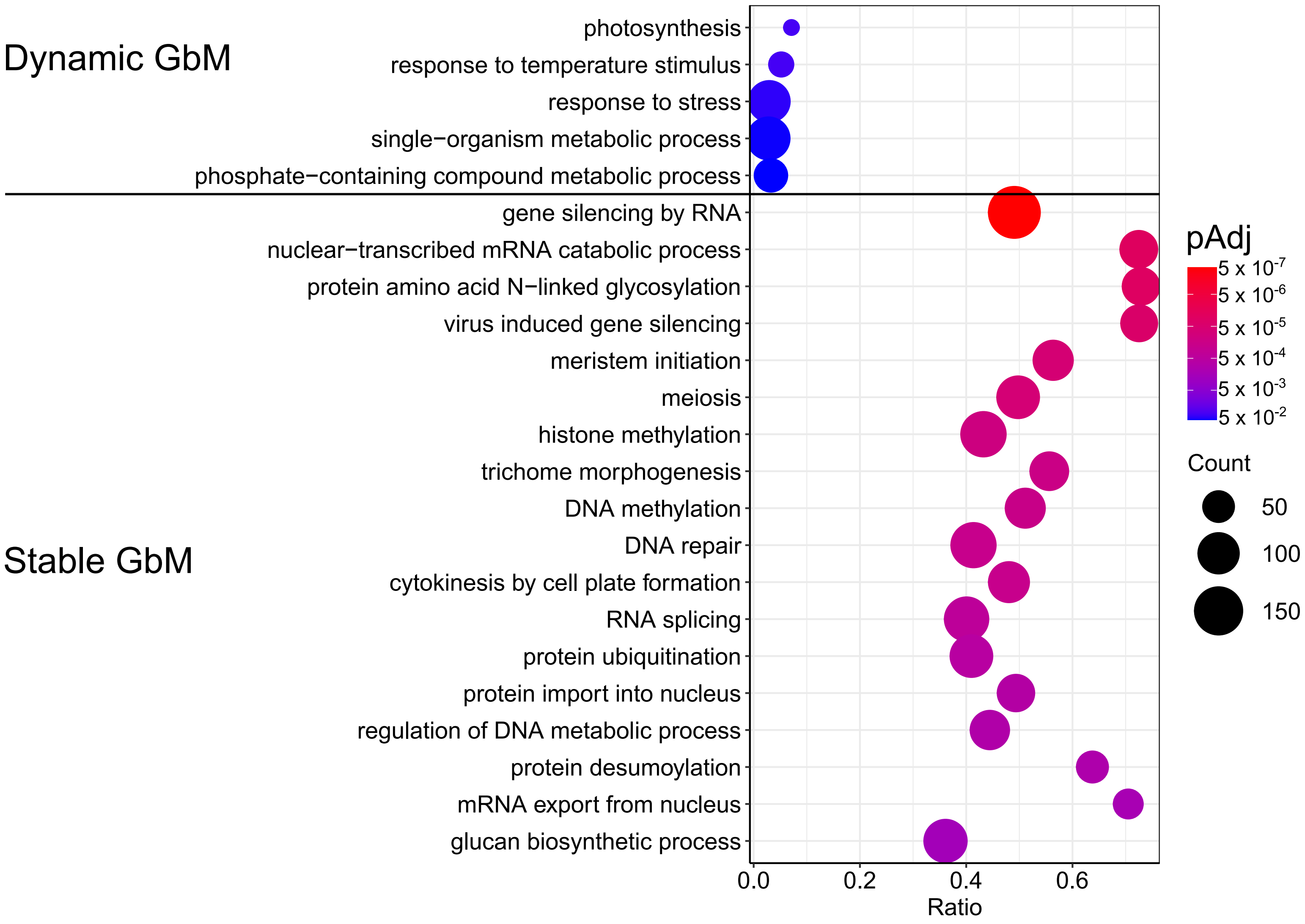


**Figure S9: Enriched GO-terms in Dynamic and Stable GbM genes**. Symbol color is a heat map scale for statistical significance and symbol size is proportional to the number of genes for the GO term. The x-axis indicates the enrichment ratio for each GbM category relative to all genes with the GO term.

**Table S1: A list of all external datasets analyzed in this study**

| Original Study | Publication doi | Raw data accession |
|---|---|---|
| Bewick, A.J. et al, (2016) PNAS. 113 (32) 9111-9116 | 10.1073/pnas.1604666113 | NCBI GEO GSE75071 |
| Bourguet P. et al, (2021) Nat. Commun. 12:2683 | 10.1038/s41467-021-22993-5 | NCBI GEO GSE146948 |
| Chen, L.Q. et al, (2017) Plant J. 174(3):1795-1806 | 10.1104/pp.16.01944 | NCBI GEO GSE73972 |
| Gutzat, R. et al, (2020) EMBO J. 39:e103667 | 10.15252/embj.2019103667 | EMBL EBI E-MTAB-5478 & E-MTAB-5479 |
| Kawakatsu T., et al (2016). Nat Plants. 2:16058 | 10.1038/nplants.2016.58 | NCBI GEO GSE79710 |
| Kawakatsu, T. et al, (2016) Cell. 166(2):492-505 | 10.1016/j.cell.2016.06.044 | NCBI GEO GSE43857 |
| Khouider, S. et al, (2021) Nat. Commun. 12:420 | 10.1038/s41467-020-20606-1 | NCBI GEO GSE141154 |
| Liang, W. et al, (2022) New Phytol. 233:722–737 | doi.org/10.1111/nph.17804 | NCBI SRA PRJNA686693 |
| Liu, Y. et al, (2018) Nucleic Acids. Res. 46:157-167 | doi: 10.1093/nar/gkx919 | http://systemsbiology.cau.edu.cn/chromstates |
| Long, J. et al, (2021) Science. 373:eabh0556 | 10.1126/science.abh0556 | NCBI GEO GSE161625 |
| Mergner J. et al, (2020) Nature, 579:409-14 | 10.1038/s41586-020-2094-2 | https://www.nature.com/articles/s41597-020-00678-w |
| Niederhuth, C.E. et al, (2016) Genome Biol. 17:194 | 10.1186/s13059-016-1059-0 | NCBI GEO GSE79526 |
| Rigal, M. et al, (2016) PNAS. 113 (14) E2083-E2092 | 10.1073/pnas.1600672113 | EMBL EBI PRJEB9919 |
| Seymour, D.K. et al, (2014) PloS Genet. 10: e1004785 | 10.1371/journal.pgen.1004785 | EMBL EBI PRJEB6701 |
| Shang J-Y. et al, (2022) J. Integr. Plant Biol. 64:2438-54 | 10.1111/jipb.13406 | NCBI GEO GSE206688 |
| Stroud, H. et al, (2013) Cell. 152(1-2):352-64 | 10.1016/j.cell.2012.10.054 | NCBI GEO GSE39901 & GSE38286 |
| Stroud, H. et al, (2014) Nat. Struct. Mol. Biol. 21:64-72 | 10.1038/nsmb.2735 | NCBI GEO GSE51304 |
| Williams B.P., et al (2022). Plant Cell;34:1189–206 | 10.1093/plcell/koab319 | NCBI GEO GSE191307 |

### Table S2: properties of EM-seq libraries sequenced in this study

| Sample | Tissue | Cycles | Total reads | Total reads after QC | Uniquely mapping reads after deduplication | % Reads uniquely mapping | Conversion rate |
|---|---|---|---|---|---|---|---|
| *met1-3* heterozygote | leaf | 2 x 150 bp | 60,407,565 | 57,704,567 | 30,318,827 | 52.6 | 99.998 |
| *met1-3* WT segregant, line 1 F2 | leaf | 2 x 150 bp | 59,741,235 | 56,772,575 | 32,355,357 | 57.1 | 99.9978 |
| met1-3 WT segregant, line 1 F3 | leaf | 2 x 150 bp | 53,407,468 | 53,362,670 | 22,649,572 | 42.4 | 99.999 |
| met1-3 WT segregant, line 2 F2 | leaf | 2 x 150 bp | 50,522,623 | 50,465,956 | 25,117,695 | 40 | 99.9989 |
| met1-3 WT segregant, line 2 F3 | leaf | 2 x 150 bp | 57,417,384 | 55,601,549 | 32,596,017 | 58.6 | 99.9978 |
| met1-3 WT segregant, line 3 F2 | leaf | 2 x 150 bp | 54,398,483 | 54,337,350 | 24,958,077 | 45.9 | 99.999 |
| met1-3 WT segregant, line 3 F3 | leaf | 2 x 150 bp | 55,237,611 | 55,193,429 | 26,779,936 | 48.5 | 99.9989 |
